# Cross-Paradigm fNIRS Brain Activity in Neonates across The Gambia and UK

**DOI:** 10.1101/2025.07.28.666710

**Authors:** I. Greenhalgh, B. Blanco, C. Bulgarelli, E. Mbye, E. Touray, M. Rozhko, L. Katus, N. Hayes, S. McCann, S.E. Moore, C.E. Elwell, A. Blasi, S. Lloyd-Fox, the BRIGHT Project Team

**Author notes:** Corresponding author: Isobel Greenhalgh. **Author contributions:** IG: conceptualisation, data analysis and manuscript writing; BB: data analysis and manuscript editing; CB: conceptualisation, data analysis and manuscript editing; EM: data collection; ET: data collection; MR: data collection; LK: data collection; NH: data collection; SMcC: data collection; SEM: project lead and manuscript editing; CE: project lead and manuscript editing; AB: conceptualisation, data analysis, manuscript writing and project supervision; SLF: conceptualisation, project lead and supervision and manuscript writing. The BRIGHT Project Team: Austin, T., Perapoch-Amado, M and Ghillia, G. Declarations of interest: none.

## Abstract

**Significance:** Neonates undergo rapid development, yet the examination of emerging brain markers across paradigms, cognitive domains and diverse global populations remains limited.

**Aim:** This study investigated whether brain responses at one-month-of-age could be interrogated across paradigms to offer deeper context-specific insights into neurodevelopment.

**Approach:** Functional near-infrared spectroscopy (fNIRS) was used to assess frontal and temporal brain responses during natural sleep in 181 Gambian (GM) and 58 UK infants during three auditory paradigms: Social Selectivity, Habituation and Novelty Detection (HaND) and Functional Connectivity (FC). Paradigm-level brain responses were analysed using threshold-free cluster enhancement and cross-paradigm comparisons of individual responses.

**Results:** At the group level, both GM and UK infants showed habituation but not novelty responses, higher inter-versus intra-hemispheric connectivity, stronger inter-hemispheric connectivity in temporal regions relative to frontal regions, stronger inter-regional connectivity between right temporal and left frontal regions, and UK infants also showed non-vocal > vocal selectivity.

**Conclusions:** Cross-cohort differences in the cross-paradigm analyses suggest context-specific developmental markers are evident within the first month of life and show high individual variability. Cross-paradigm analyses revealed that greater vocal selectivity (UK) was associated with higher inter-hemispheric connectivity, potentially allowing us to identify biomarkers of more mature neurodevelopment within the first weeks of postnatal life.

## 1.0 Introduction

The first 1000 days of life – from conception to two years of age - have been posited to represent those most integral to human neurodevelopment, as rapid and prolific neurogenesis, synaptogenesis, and synaptic pruning occur in response to genetic-environmental interplays^1,2^. During this time, development across social, emotional and cognitive domains is witnessed as the brain adds and prunes its connections to build functional networks, with research demonstrating that such early abilities lay the foundation for later education, workplace, health and wellbeing outcomes^4,5,6,7^. A key developmental stage is the perinatal transition to postnatal life - a period of important transition and change^8^. However, examining this period is challenging: access to newborn infants during a period of heightened vulnerability and time constraints for parents can be limited; given the age of the participant identifying and conducting research during stable infant states of alertness can be less predictable; and infant-friendly paradigms, particularly for neuroimaging research, that can be employed in early life have for a long time been less common. In recent years, technological advances in neuroimaging techniques such as functional near-infrared spectroscopy (fNIRS), electroencephalography (EEG) and functional magnetic resonance imaging (fMRI) and paradigms implemented during sleep (particularly in the auditory domain), have allowed some of these barriers to be overcome^9,10,11,12^. Subsequent research has highlighted the non-linearity of developmental trajectories across infancy, childhood and adolescence^2,7,13^. Yet we still lack objective insights into what constitutes typical and atypical neurodevelopment, especially across lower income settings, and when considering the first months of life.

In low-and middle-income countries (LMICs) it has been estimated that around 30% of children fail to meet their developmental milestones^14^. Exposure to adversity, including undernutrition, poor parental mental health, infection and overcrowding, is more common in such settings, compared to higher income countries (HICs), and such exposures early in life associated with poorer developmental outcomes across social and cognitive domains^5,14,15,16^. Understanding of the effects of early exposures, and possible buffers, across contexts is critical.

The Brain Imaging for Global Health (BRIGHT) project aimed to fill this essential gap in early neurodevelopmental research, through a detailed examination of infant neurocognitive, social and functional brain development from the first weeks of postnatal life until five years of age, across a higher income cohort in the UK, and an under-researched, low income cohort in The Gambia^17^. A battery of neuroimaging paradigms (EEG and fNIRS) were conducted from one-month of age, in order to identify and map longitudinal neurodevelopment across settings. Thus far, findings from the BRIGHT project have helped the research community to better understand functional brain connectivity, and neurocognitive and social trajectories, from five months to five years^18,19,20,21,22^. To date, findings suggest that the period of postnatal development between 0 – 6 months may be significantly impacted by exposure to risk factors^20,23^. However, the study of brain functionality at the earliest time point at one-month-of-age in the BRIGHT project has yet to be fully explored. Given that we know a range of social and neurocognitive skills have their onset in early postnatal life and indeed during pregnancy, further research into this early timepoint is required^8^.

Infants are born into a social world. The capacity to engage with others, to differentiate between social and non-social stimuli, and to infer information through language and non-verbal communicative gestures, is part of what makes us human, and such skills are utilized in everyday life. While typically, the emergence of social (over non-social) stimuli was posited to characterize early infant development across the first years of life, increasing research which examines the very first months post birth indicates that this selectivity can be particularly context/stimulus dependent. For example, while we see evidence of visual social selectivity from birth^24^, in the auditory domain this selectivity may be more stimulus specific and variable across individuals. At the group level this results in non-social selectivity (to environmental non-vocal sounds) predominating in the first weeks, before transitioning towards social selectivity (to vocalisations) around 4-8 months of age^10,25^. Such non-social selectivity has also been reported to be more widespread across several brain regions, while emerging social selectivity from 4 months and beyond appears to be more localised to anterior temporal and inferior frontal cortices^19,25,26,27,28^.

Social and cognitive paradigms often interrelate: focusing on social stimuli requires attentional skills, while holding, updating and responding appropriately to social stimuli can draw upon working memory and inhibition^13,19,29,30^. Notably, neurocognition, for example processes such as reasoning, attention and memory manifest at both the brain and behavioural level, has been identified as one of the best indicators of later neurodevelopment^31^, especially in predicting development in self-regulation and executive functions^32^. Given the critical nature, and interdependency of both social and cognitive abilities throughout life, research which seeks to elucidate the development of such functions in parallel, is needed.

Two early neurocognitive skills which can be assessed in early infancy, and which map onto a myriad of later outcomes, are the tethered abilities of habituation and novelty detection (HaND). The ability to decrease neural and behavioural responses to inconsequential, repetitive stimuli (habituation), and to recover the response when new stimuli are encountered (novelty detection) has been associated with measures of brain efficiency, cognition, education, and IQ, with improved HaND abilities conferring improvements across these developmental domains^33,34,35,36,37,38^. During infancy, differences in both habituation and novelty detection (HaND) have been cited across Gambian and UK infants, with both displaying attenuated responses to repeated stimuli at 5-and 8-months of age, while novelty detection also emerges at this time point in UK, but not Gambian, infants^19,22^. In contrast, Gambian infants continue to display habituation responses, with novelty responses emerging at 18-months of age, while UK infants demonstrated attenuation of both habituation and novelty recovery, such that there were no significant responses by 18-months of age^22^. Of interest, the BRIGHT project examined HaND responses across the first years of life in Gambian infants, reporting modest correlations in habituation responsess across both EEG and fNIRS across one and five months, but not at 18-months, while novelty responses correlated at 5-and 18-months only^11^. Such findings demonstrate the complexity of early neurodevelopmental trajectories both across and within diverse cohorts and settings.

While task-specific paradigms play an integral role in helping understand the emergence of specific skills and their localisation, research examining the networks that underpin such skills is also essential. The foundations of many functional networks develop during pregnancy.

However, postnatally, huge shifts occur as infants transition from the enclosed, muted and darker environment of the womb to a loud, bright and stimulating environment. As infants increase in age, segregation and integration between, and within, certain neural networks reflects both cognitive and social-emotional gains^7,39^. For example, functional connectivity of fronto-temporal networks has been reported to predict infant age and brain maturation, while frontal inter-hemispheric connectivity at five-months has been found to predict cognitive ability at pre-school age^20,40^. Similarly, inter-hemispheric connectivity of sensorimotor networks increases over the first month of life, as infants are exposed to more perceptual stimuli^41^. During this time, research supports the idea that increases in long-range, inter-hemispheric connections, and decreases of shorter-range, intra-hemispheric connections, due to synaptic pruning, support the transition to the postnatal world and scaffold the development of both early and downstream functions^20,42,43^. Yet variability in such functional development can be seen across settings, with exposure to socioeconomic inequalities stronger predictors of brain dynamics than age or cognition in a large, diverse study of healthy adults across many lower and higher income settings^44^. In early infancy, functional connectivity can be disrupted by exposure to adversity, predicting poorer cognitive outcomes in childhood^20^. Whether such disruptions manifests differently across settings in the first month of life is unknown.

Given the concurrent and predictive interplay of both cognitive and social development, studies which examine the neural correlates of these skills in tandem, while also considering underlying functional networks, can provide more in-depth insights into early neurodevelopmental markers and how such skills may interrelate both at the paradigm and network level^4,45^. By including and assessing multiple indicators of early brain development, more robusts insights may be gained into typical developmental profiles across settings, and may help to decipher whether they appear to draw upon overlapping or distinct mechanisms. To date, no research has examined the foundational stages, and interrelated development of, neurocognitive and social skills and associated functional brain connectivity, in the first weeks of life, across two diverse, real-world settings.

The present study sought to address this by (i) examining early Gambian and UK infant neurodevelopment at the relatively under-researched age point of one-month, and (ii) examining whether brain responses across neurocognitive and social domains are inter-related, and how they more widely relate to functional brain connectivity. Data from a *Social Selectivity* paradigm, *Habituation and Novelty Detection (HaND)* paradigm, and *Functional Connectivity* paradigm were collated and examined both at the paradigm-level, and in cross-paradigm analyses. This data was collected in a single session, during sleep at one-month in infants using functional near-infrared spectroscopy (fNIRS). Much of this work was exploratory to try to understand whether cross-paradigm markers of infant development could be identified both within and across settings, and whether these could be used to group infants according to their developmental maturity.

However, the following hypotheses were put forward: (1) infants with more mature connectivity profiles (i.e. stronger overall inter-hemispheric connectivity) will show more rapid specialisation of social discrimination responses (i.e. will display increased selectivity towards vocal (social stimuli) compared with non-vocal (non-social) stimuli) as well as more specialised information processing (more robust habituation and novelty detection responses); and (2) if the social and HaND paradigms are actually interrogating similar perceptive abilities (due to the use of social human stimuli in both) or reliant on similar underlying cognitive processes infants with stronger vocal (social) responses will have more robust habituation and novelty detection responses. If however, individual responses across these paradigms are unrelated this would suggest that they are each able to identify a unique biomarker of early development.

## 2.0 Methods

### 2.1. Participants

The BRIGHT project recruited participants across two study sites (for full details of the participating families see^17)^:

The Gambia (GM) Cohort: Participants were recruited during pregnancy from the village of Keneba and neighbouring villages in the West Kiang District. Prospective participants were approached and provided with information pertaining to the project. Should they wish to take part, written or thumbprint consent was obtained from participants, with the latter method employed for participants who could not write (the study was approved by the joint Gambia Government-MRCG Ethics committee (ref # SCC 1451). Of the 204 participants enrolled in the BRIGHT project at the one month time point, 181 undertook the fNIRS session at one-month-of-age, the inclusion rates for each paradigm are outlined in the Results section.

The Gambia is situated in sub-Saharan Africa and it is the smallest country on the continent, with a relatively young population (45% is age 0-14, while 4% is 65 or older)^46^. Families typically live in larger family units. A majority of the population (particularly women) have not completed higher education and a majority of rural dwellers work in agriculture, mainly for subsistence^47^.

There is a high prevalence of growth faltering in children, which is notably affected by seasonally-impacted food insecurity driven by weather patterns characterized by two highly differentiated seasons (dry and rainy, each spanning half a year) that modulate the availability of key nutrients^48,49^. For a detailed characterization of the Gambian population, see^46^. Ethnicity in the West Kiang District of The Gambia is predominantly Mandinka (79.9% of the population^49^) with its unique language and cultural characteristics. To avoid confounds caused by multiple translations of the project protocols, GM cohort families recruited into the project were predominantly of Mandinka ethnicity (i.e. the main language spoken at home was Mandinka).

The UK Cohort: Participants were recruited during pregnancy in the Rosie Hospital, Cambridge University Hospitals, Cambridge, UK, between 32 to 36 weeks’ gestation. Upon approach mothers were given information sheets detailing the BRIGHT project and were then contacted via email or phone at a later date to determine whether or not they wished to participate. If participants did wish to partake in the BRIGHT project, informed consent was gathered using written forms, in alignment with ethical approval (National Research Ethics Service Committee East of England (REC reference 13/EE/02000). Of the 61 families enrolled in the project at 1 month-of-age, 58 undertook the fNIRS session at one-month-of-age, the inclusion rates for each paradigm are outlined in the Results section.

The majority of participants within this cohort lived within the city of Cambridge or within urban or rural communities within a 20-mile radius. Demographically, the population in Cambridgeshire is representative of that across the UK with regard to ethnicity, employment rates and family structure^50^. The area however differs from the rest of the UK with regard to levels of education within the population, with twice as many inhabitants holding a higher education degree^50^.

Prerequisites for inclusion in the BRIGHT project were (1) that infants were carried to full term (37-42 weeks gestation), and (2) for UK participants only, they had a normal birth weight (>2.5kg). The latter constraint was not placed upon participants in the GM cohort, given the higher rates of growth restriction due to dietary deficiencies, contamination and infection^19^.

However, Gambian infants who had experienced severe growth faltering (weight-for-height or head circumference z-score less than-3 according to WHO standards) were excluded.

### 2.2 fNIRS Data Acquisition

Infants wore a custom-designed fNIRS cap (NTS optical topography system, Gowerlabs Ltd, UK) which consisted of two arrays of 9 channels each, that covered left and right hemispheres (Figure 1 below). The system used two continuous wavelengths of near-infrared light (780 and 850nm) and a sampling frequency of 10Hz. Arrays were designed to cover frontal and temporal regions, with source-detector pairs positioned at a 2cm distance.

**Figure 1.**
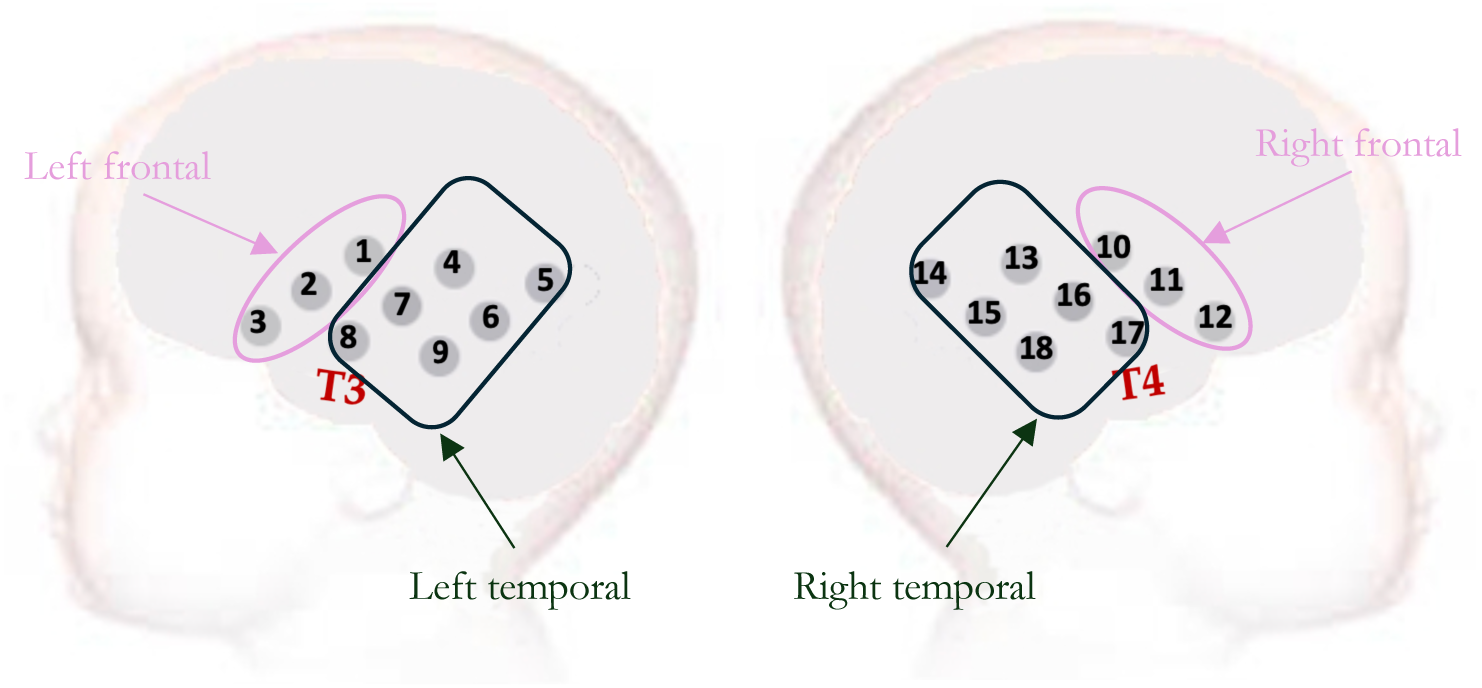
Layout of the 18-channel arrays used at 1-month, covering frontal and temporal areas with the 10-20 T3/T4 positions identified. The region-of-interest channel clusters used for averaging in Functional Connectivity analyses are also indicated here.

Before placing the cap on the infants, head measurements were taken including head circumference, pre-auricular to pre-auricular over the head and across the forehead, and nasion to inion. Photographs were then taken, which, combined with head measurements, allowed for later registration and estimation of the cortical areas relating to each channel location.

Subsequently the headgear was placed and secured with an additional band to ensure proper fitting and prevent slipping during the session. If the infant was awake during this process, researchers then waited for the infant to fall asleep before starting the data acquisition. At the start of the session, the infant was held, swaddled, by either the parent or researcher, and allowed to fall asleep naturally in their arms. Research sessions occurred in the daytime during the infant’s usual naptime to try to ensure success with sleeping. The infant was video-recorded to allow the researcher to check the position of the infant across the session, and monitor sleep stages/arousal. Infants were asleep in their parent’s or the researchers’ arms or placed on their laps, and positioned approximately at a distance of one metre from the screen and the speakers that played the sounds. During the data collection session, stimuli were presented via Logitech Z130 speakers which were connected to a laptop on which the stimuli script was played using MATLAB, Task Engine (Task Engine, sites.google.com/site/taskenginedoc) and Psychtoolbox^51,52,53^. The sound volume was adjusted to an average of 60dB when the sound reached the position of the infant’s head.

Once the infants were sleeping the paradigms were presented in the same order (Figure 2 below), first the *Social Selectivity* Paradigm, then the *Habituation and Novelty Detection (HaND)* Paradigm and finally the *Functional Connectivity* Paradigm. If the infant became fussy or began to wake the session was paused to allow them to settle again. If they awoke, did not return to sleep and were calm then the session continued, however if they became upset or excessively fussy the session was ended. Therefore, the validity of data is highest for the Social Selectivity data as this paradigm was conducted first. In total the data collection lasted for approximately 30 minutes.

**Figure 2.**
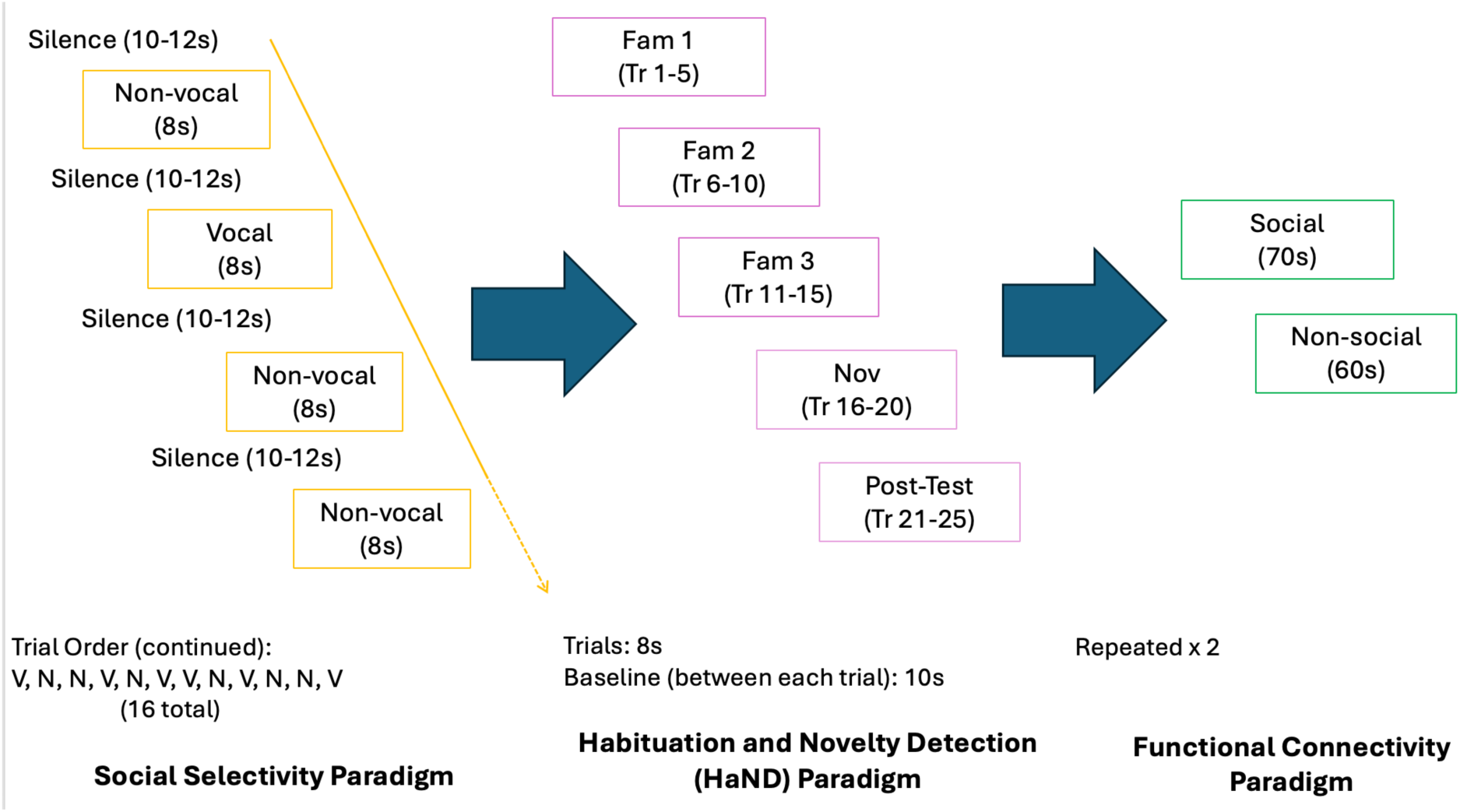
Order of paradigm presentation at one-month in the BRIGHT project. On the left the Social Selectivity paradigm can be seen to be made up of 16 trials, with pseudo-randomised order of vocal (V) and non-vocal(N) trials. After, (in the middle), the HaND paradigm was undertaken, comprised of Familiarisation 1 (Fam1 – trials (Tr) 1-5), Familiarisation 2 (Fam 2 – trials 6-10), Familiarisation 3 (Fam 3 – trials 11-15), Novelty (Nov – trials 16-20, which comprised a change in speaker), Post-Test (trials 21-25, back to original speaker). Finally, Functional Connectivity (FC – right) was undertaken with twice repeated social (singing nursery rhymes – 70s) and non-social (auditory sounds of toys – 60s) audio recordings.

### 2.3 Stimuli and Design

#### 2.3.1 Social Selectivity Paradigm

While at later age points in the BRIGHT project^21^, infants were exposed to visual and auditory stimuli in the *Social Selectivity* paradigm, at one month of age the paradigm only contained auditory stimuli given this was undertaken during sleep^10^. Auditory stimuli consisted of both vocal (social) and nonvocal (nonsocial) sounds, each of which lasted 0.37-2.92 seconds and which were presented in clusters of 8 seconds. Vocal stimuli were comprised of non-speech but vocal human sounds (laughter, crying, yawning and coughing), while non-vocal stimuli were environmental non-social sounds (running water, bells, and rattles). Each presentation of vocal or non-vocal sounds contained four different sounds within that condition, with a baseline preceding each experimental trial. Both vocal and non-vocal stimuli averaged the same sound intensity and duration. Further information on the *Social Selectivity* paradigm stimuli can be found in^10,54^. The condition trials were presented in pseudo-random order (N (non-vocal), V (vocal), V, N, V, N, N, V, N, V, V, N, V, N, N, V) with a jittered length 10 to 12 sec silent baseline period directly preceding and then after each consecutive trial, with a maximum number of 16 condition trials presented.

#### 2.3.2 Habituation and Novelty Detection (HaND) Paradigm

The HaND auditory stimuli consisted of spoken sentences in either English (UK: “Hi baby! How are you? Are you having fun? Thank you for coming to see us today. We’re very happy to see you”) or Mandinka (The Gambia: “Denano a be nyadii. I be kongtan-rin? Abaraka bake elan aa kanan njibee bee, n kntanta bake le ke jeh”). Sentences lasted 8-seconds in length, with one sentence counted as one trial, and each trial preceded by a 10s silent baseline. During the habituation phase, the sentence was repeated 15 times by a female speaker. During the novelty phase, the speaker was then changed to a male speaker for five trials, before a final five re-familiarisation trials by the original female speaker. This resulted in a total of 25 trials. The stimuli were recorded at 48Khz sampling rate before being edited via Audacity software v2.2.1 to normalise to a peak amplitude of-1dB SPL. Stimuli were also converted to mono from stereo.

To better capture the response signature of the *HaND* paradigm, trials were further grouped in epochs of five trials each, using the same methodology as published analyses of this protocol^19^. Then, epochs were defined as follows: Familiarisation 1, from now on labelled as Fam 1, included trials 1 to 5; Fam 2, with trials 6-10; Fam 3, with trials 11-15; Novelty, from now on labelled Nov, with trials 16-20; re-familiarisation, labelled here re-fam, included trials 21-25.

#### 2.3.4 Functional Connectivity

After the *Social Selectivity* and *HaND* paradigms were complete, if the infant was still asleep, functional connectivity data were recorded using the paradigm implemented at older age points^20^. The protocol consisted of playing videos of UK or Gambian (appropriate for each population) male and female adults signing and singing nursery rhymes (70 sec) and of toys in action (such as rattles and spinning bells/mirrors) (60 sec). The sequence was repeated twice, with a maximum recording time of 260 sec. For the majority of infants this was performed while they were sleeping and therefore only the auditory stimuli would have been available. In cases when infants were awake but quietly resting, the screen that displayed the stimuli was not in direct eyesight of the infant, to ensure this was also an auditory-only paradigm.

### 2.4 fNIRS Pre-Processing and Analysis: Social Selectivity and HaND Paradigns

Post data collection, the optical density datasets were inspected visually for excessive noise and to ensure all event markers were present. Next, channels with poor quality data were flagged for exclusion in subsequent analysis using QT-NIRS (https://github.com/lpollonini/qt-nirs) with cut-off frequencies adjusted to capture infant heart rate [1.5,3.5] Hz, Scalp Coupling Index (SCI) = 0.7, and Peak Spectral Power (PSP) = 0.1; channels that did not meet the SCI and PSP thresholds for at least 70% of their temporal window were flagged. Datasets with more than 40% flagged channels (more than 7 channels) were excluded from further analysis. The remaining datasets were pre-processed following guidelines for infant research^55,56,57^ with Homer2 V2 with a pipeline that included: Spline interpolation (p = 0.99); wavelet denoising (inter quartile range (IQR) = 0.8), and low-pass filtering, with a cut-off frequency of 0.6Hz; and conversion from optical density to oxy-and deoxy-haemoglobin concentration using the Beer-Lambert Law^58^ with wavelength-dependent differential pathelength factors (DPFs) of 5.22 and 4.23^55^. After pre-processing, only datasets with a minimum number of 3 trials for the two conditions of the *Social Selectivity* paradigm, and only datasets with a minimum of three trials in the first and last familiarisation epochs (Fam1 and Fam3) of the HaND paradigm continued to be considered for further analysis.

In order to understand whether significant differences in activation could be seen across conditions, channels, and the time course of the hemodynamic response within each paradigm, a threshold free cluster enhancement (TFCE) approach was undertaken^59^. Threshold free cluster enhancement offers benefits compared to other statistical methods given it takes a data driven approach but also its calculation of spatial clusters based off distance between channels (defined as 20mm in the current study) in contrast to other methods such as cluster permutation analysis, which require subjective specification of channels per cluster. TFCE uses statistical analyses performed on individual hemodynamic responses for generating a channel-wise output that accounts for both the height (or amplitude) of the response and the extent (or spatial and temporal characteristics), with neighbouring values therefore contributing to the new value per channel. These are then developed into p-values, corrected for multiple comparisons (using the Benjamini-Hochberg correction^60^; False Discovery Rate (FDR)), which indicate the significance of each cluster and can further infer information about time windows during which significant responses are witnessed across the whole temporal haemodynamic response. If a response was found to be significant for under 1 second, this was considered to not be robust and did not go forward into further analyses. In the present study TFCE was used to both examine group level outcomes within each paradigm and to identify the time windows and regions of interest to be used for data extraction for cross-paradigm analyses. Time-windows are calculated using the median, given the prevalence of artifacts and outliers in fNIRS data, ensuring chosen time windows are more robust to such deviations from true activation.

TFCE was run for both condition versus baseline and condition vs condition contrasts. For the *Social Selectivity* paradigm these included: (1) non-vocal condition (N) > baseline; (2) vocal condition (V) > baseline; (3) N vs V contrast. Then, in channels with a significant N > V contrast, a significant N > baseline was required to consider the N > V result as valid; and in channels with significant V > N, a significant V (> baseline) was required.

For the *HaND* paradigm, tests included: (1) Fam1 > baseline; (2) Fam3 > baseline; (3) Novel > baseline; (4) Fam1 vs Fam3 contrast (to obtain a Habituation response); and (5) Novel vs Fam3 contrast (to obtain a Novelty detection response). Following the same principle as in the Social paradigm, to consider significant habituation (Fam1 > Fam3), a significant Fam1 (> baseline) was required; and to consider significant novelty detection (Novel > Fam3), a significant Novel (>baseline) was required.

### 2.5 fNIRS Pre-Processing and Analysis: Functional Connectivity

Data preprocessing and analysis was carried out in MATLAB (MathWorks, Natick, MA) following the same procedure as found in^20^. First, channel quality was assessed using QT-NIRS (https://github.com/lpollonini/qt-nirs), with the same parameter settings as for the Social and HaND data. The raw intensity data was then converted to optical density (OD) in Homer2 before undergoing bandpass filtering (0.009 – 3 Hz). The mean of signals across the array were also regressed out using the global signal regression (GSR), to reduce physiological noise that can infiltrate fNIRS data^61,62^. OD data was then passed through a second bandpass filter (0.009 – 0.08 Hz), before motion artifact rejection. This excluded 5 seconds before and after the artifact occurred, and made use of the global variance of temporal derivatives (GVTD, STD=5) in the NeuroDOT toolbox (https://www.nitrc.org/projects/neurodot/)^63^. Only infants with a minimum of 120 seconds of valid FC data post pre-processing were included in further analysis. Finally, OD data was converted into HbO and HbR concentrations as above (Delphy et al., 1988).

To reduce the number of multiple comparisons incurred at channel level, the arrays were partitioned in two regions per hemisphere, as illustrated in Figure 1, resulting in four regions in total. The sections were grouped into frontal and temporal regions (for a total of four regions) based upon previous co-registration of the BRIGHT fNIRS arrays^64^. Average HbO and HbR concentration changes across all channels within each ROI (left frontal, left temporal, right frontal and right temporal) were used to explore connectivity between the regions. For each participant, the Pearson-r correlation values between all the ROIs was calculated for both HbO and HbR, resulting in a 4×4 matrix of section-pair correlations. Correlation coefficients were computed for each intra-and inter-hemispheric combination, then Fisher-z transformed. Overall inter-hemispheric and intra-hemispheric connectivity scores were calculated by averaging the 4 inter-and the 6 intra-hemispheric connectivity measures for each participant. Total intrahemispheric and interhemispheric connectivity was also calculated by summing the total between-region and within-region correlations.

### 2.5 Cross-Paradigm Analyses

Once ROI’s and time windows driving significant activation had been identified for each paradigm, these were then used to guide data extraction for cross-paradigm analysis. These analyses focused on the HbO results. HbR results can be found in Supplementary analyses. For the *HaND* paradigm, four-second time windows were calculated based upon significant activation in Fam 1, with common channels active in both the UK and the GM cohorts for Habituation (Fam 1 > Fam 3) extracted. Given the similarity of the ROI and time window where the strongest haemodynamic response was evident across the two cohorts (see results), this was kept consistent for the cross-paradigm so that responses common across cohorts could be examined. For the *Social Selectivity* paradigm, no activation was found at the condition contrast level in the GM cohort (see Results). Therefore, the UK cohort selectivity (N > V) ROI was identified and used across cohorts, in keeping with^54^, to decipher whether individual variability couild still be picked up in cross-paradigm analyses in the GM cohort despite lack of significance at the group level. However, given that the time window for the peak of the haemodynamic responses for both Nonvocal and Vocal conditions did not overlap between the two cohorts, two different time windows were selected, as outlined in the results.

Finally, for the *Functional Connectivity* paradigm, ROI’s were pre-defined in line with other publications of this data^20^ (Figure 1). The type of connectivity subsequently identified to be the strongest drivers (interhemispheric versus intrahemispheric), were then taken forwards to be used in cross-paradigm analyses, post calculation of Fisher’s transformed z-scores. Subsequently, Pearson coefficient correlations and t-tests were performed to examine the relationships between cross-paradigm brain markers.

## 3.0 Results

As shown in Figure 3, in the GM cohort, of the 204 infants enrolled in the project at one month-of-age, 181 infants had fNIRS data collected at one-month for the *Social Selectivity* paradigm, 180 for the *HaND* paradigm and 175 for the FC paradigm. Note that three participants were withdrawn from the project due to a diagnosis of a developmental delay, and the majority of the 19 participants who missed a visit did so due to living away from their main family home at the time of the session (due to giving birth elsewhere) and thus being unable to travel to MRC Keneba. Following data quality control checks and data pre-processing steps for valid channel and trial data 148 had viable social data (81.8% valid), 136 had viable HaND data (75.6% valid) and 122 had FC data (69.7% valid). In the UK cohort, of the 61 infants enrolled in the project at one month-of-age, 58 infants had fNIRS data collected at the one-month visit (56 for the FC paradigm). Following data quality control checks for infant fuss out, experimental error in script or headgear placement and the data pre-processing steps for valid channel and trial data 46 infants had viable Social data (79.3% valid), 38 had viable HaND data (65.5% valid) and 41 had valid FC data (58.9% valid).

**Figure 3:**
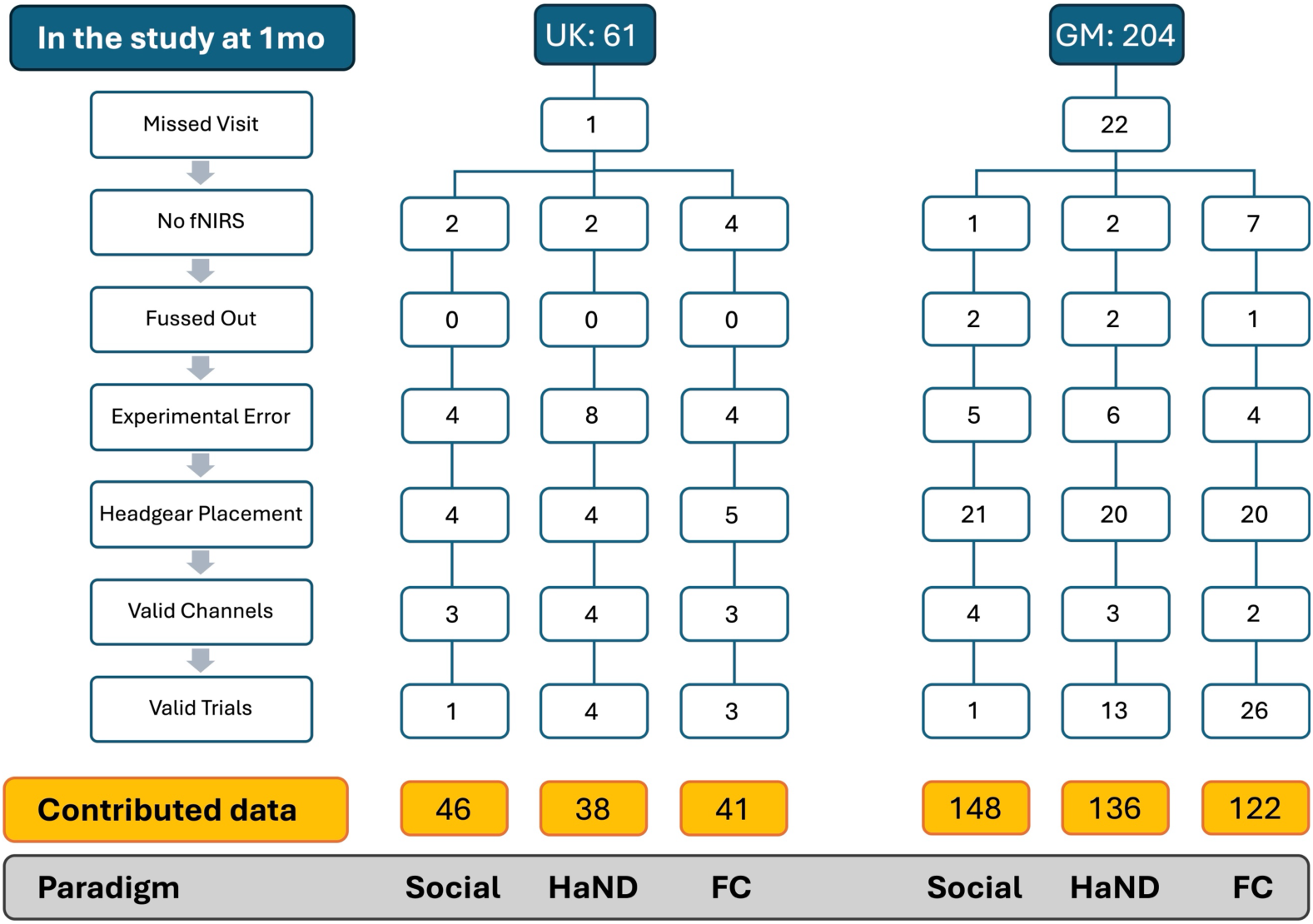
Reasons for exclusion of participants from the fNIRS analysis from the Gambian (GM) and UK cohorts across the three paradigms: Social - Non-Social (Social), Habitutation and Novelty Detection (HaND) and Functional Connectivity (FC). The reasons for exclusion included: No Session - participant did not attend the 1 month session; No fNIRS - participant attended the session, but fNIRS data was not collected for all or one of the paradigms; Fussed Out - infant woke up and cried during scanning so the recording had to be terminated; Experimental Error - collected data is invalid due to missing photos of headgear placement, missing event markers in the data collected or technical issues; Headgear Placement - collected data is invalid due to wrong placement of the fNIRS headgear or a pilot version of the headgear; Valid Channels - collected data is invalid due to too few valid channels; Valid Trials - collected data is invalid due to too few valid trials.

For the cross paradigm analyses, infants with valid data across all three paradigms totalled 30 in the UK cohort (mean age - 33.7 days (SD = 6.232); mean gestational age at birth – 38.7 weeks (SD = 3.857); 14 female / 16 male) and 107 in the GM cohort (mean age – 36.6 days (SD = 6.336; mean gestational age at birth – 38.7 weeks (SD = 1.254); 56 female / 51 male).

### 3.1 Within Paradigm-Results

#### 3.1.1 *Social Selectivity* paradigm GM Cohort

For HbO, TFCE analyses revealed left hemispheric temporal clusters with non-vocal versus baseline (N) significance ( channels 4, 5, 6 and 9) (Figure 4 below). This was driven by activation from 8.25-16.75 seconds post-stimulus onset (PST). In the right hemisphere, medial-posterior clusters drove HbO activity (channels 15, 16 and 18) from 9 – 16 seconds PST. For HbR, overlap was found in the left hemisphere, with temporal clusters driving N versus baseline significance (channels 4, 5, 6, 7 and 9) from 10.5 – 17.25 seconds PST. In the right hemisphere, frontal and temporal clusters drove significance (channels 10, 13, 14, 15 and 18) from 10.25 - 17 seconds PST. For the vocal versus baseline contrast (V), widespread activity across frontal and temporal regions drove significant right hemispheric HbO activation from 6.5 – 16 seconds (channels 10, 13, 15, 16 and 18), but no activation was found in the left hemisphere. Similarly, widespread HbR activation from 10 – 17 seconds PST drove activation in the right hemisphere (channels 10, 13, 15, 16 and 18), while activation in the left hemisphere was witnessed across a temporal cluster (channels 6, 7 and 9) from 13.75 – 17 seconds PST.

**Figure 4.**
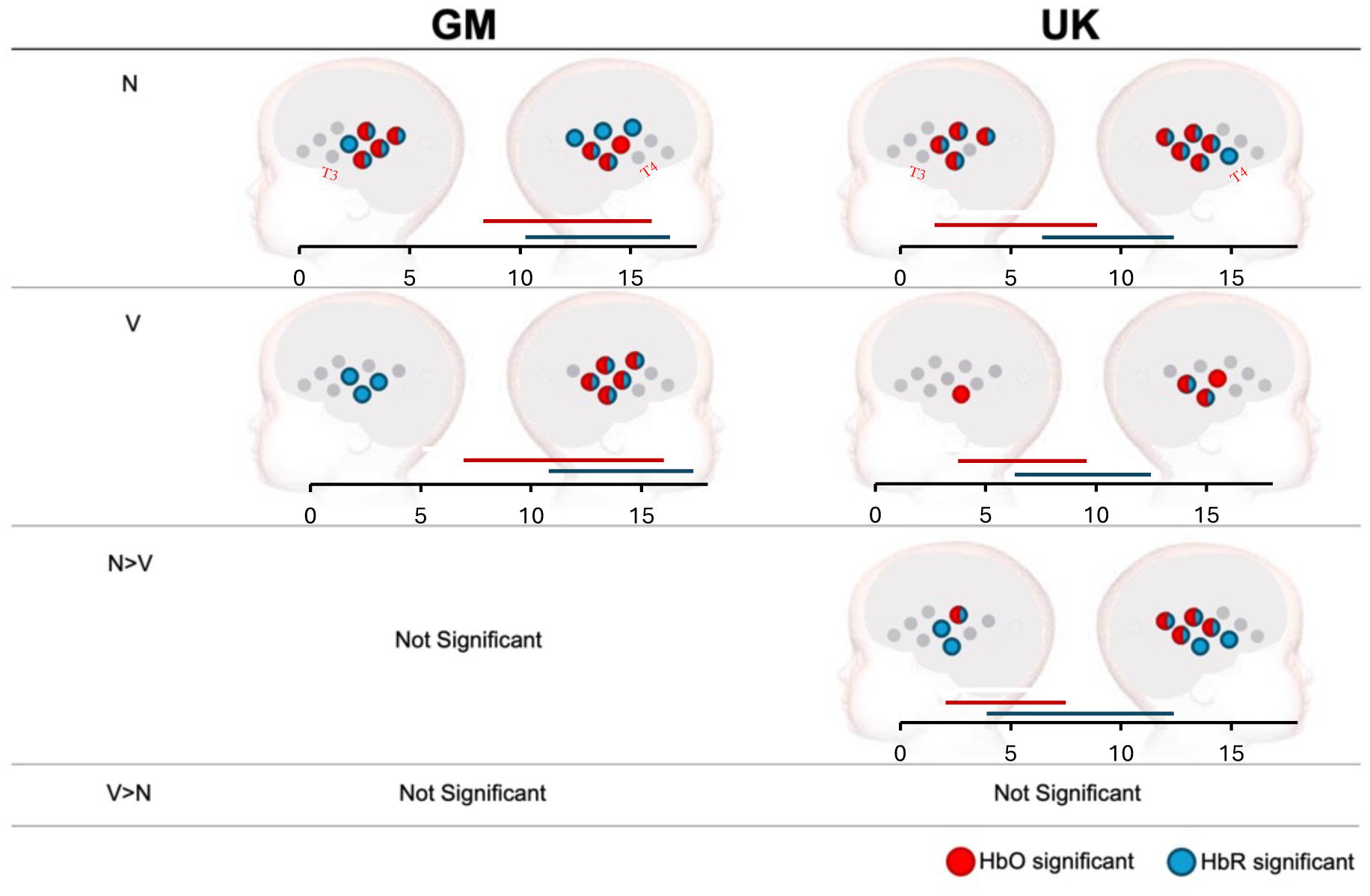
Significant channel clusters driving significant activation in the Social Selectivity paradigm across GM (left) and UK (right) cohorts. Red indicates channels with significant HbO, blue those with significant HbR. Bars displaying significant time (in sec from stimulus onset driving activation can be found below each plot. For the GM cohort, data using the UK ROI N>V was extracted for the time window of 9-13 seconds (HbO) and 10-14 (HbR). For the UK cohort, this same ROI was used but with time windows 5-9 seconds (HbO), and 7-11 seconds (HbR).

When contrasting non-vocal and vocal responses directly, no significance was found in either direction (N>V,V>N). A summary of these results can be found in Figure 4 below.

Overall, median time windows driving N and V significant activity were 6.5 – 16 seconds post-stimulus onset for HbO, and 12.25 – 17 seconds post-stimulus onset for HbR. In TFCE all significance reported was at the p<0.025 level. Subsequently, four-second time windows were used to extract data for cross-paradigm analyses. Given the wide time windows identified for HbO and HbR, four-second windows that fell into the middle of these were used: 9-13 seconds post-stimulus onset for HbO and 13-17 seconds post-stimulus onset for HbR. Given in the GM cohort no significant contrasts were found for N>V or V>N, the ROI identified for N>V in the UK was taken, as below, to explore individual variability in these responses despite the lack of group-level significance. Furthermore, as shown in Figure 5, the group grand averaged HRF across all channels found to show significance for each condition would suggest that overall the response to N was larger than V in the GM cohort.

**Figure 5.**
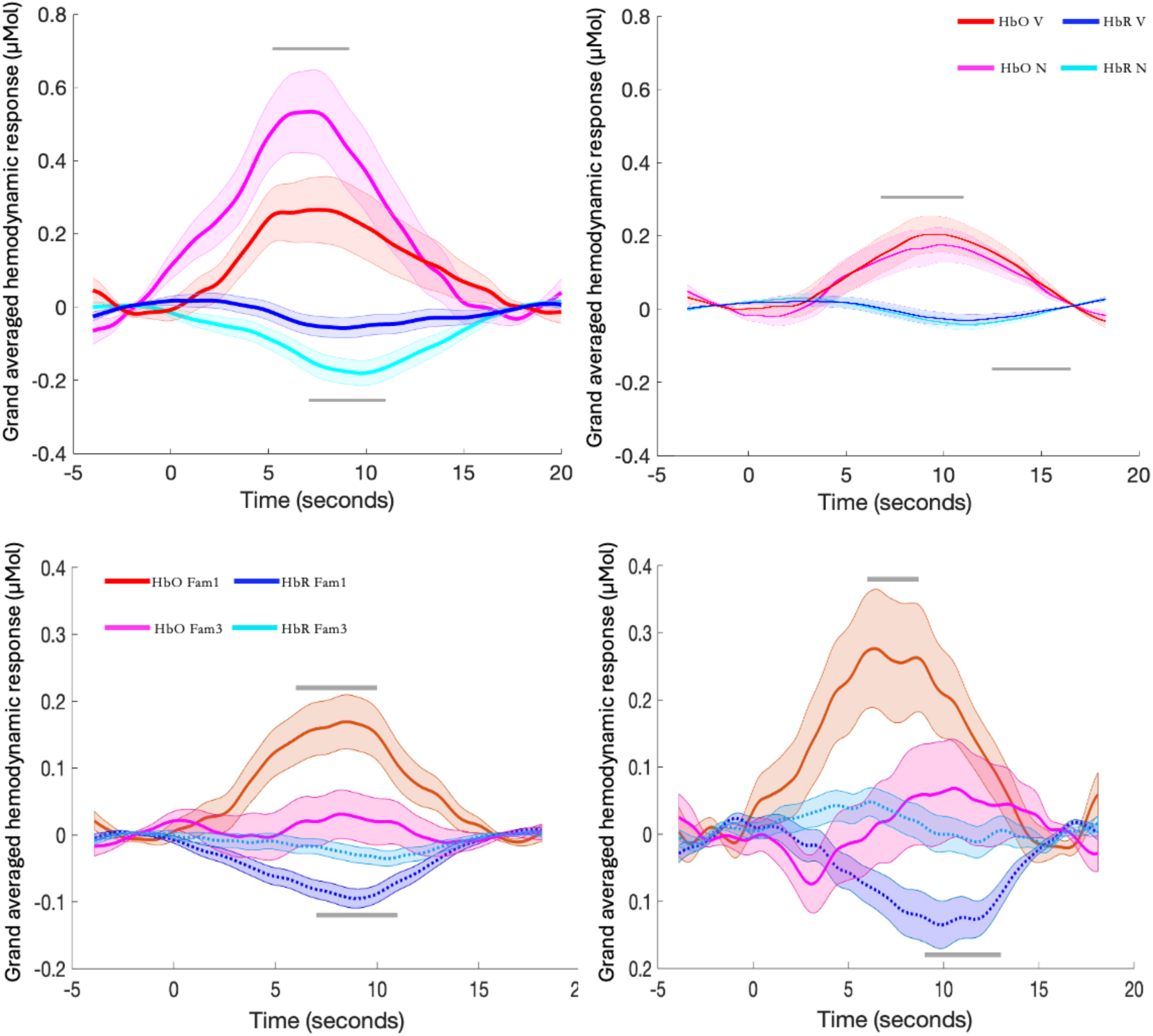
Group averaged time course (mean ± SEM) across channels found to display significance in each of the functional paradigms for the GM (left) and the UK (right) cohorts. Stimulus onset occurs at Time – 0 seconds. Grey lines indicate the time window for which mean signal change was calculated. The top row shows responses for the Social Selectivity paradigm; each plot displays HbO V (vocal – red) and HbO N (non-vocal - magenta) as well as HbR V (blue) and HbR N (cyan). The lower row displays responses for the HaND paradigm; each plot displays HbO Fam 1 (red) and HbO Fam 3 (magenta) as well as HbR Fam 1 (blue) and HbR Fam 3 (cyan) across channels found to display significance in Fam1 or Fam3 conditions.

### UK Cohort

For HbO similar to the GM cohort, the TFCE analysis revealed widespread activation in right hemispheric temporal clusters for N versus baseline contrasts (channels 13 – 16, 18) from stimulus onset to 10 seconds post. Similarly to the GM cohort, in the left hemisphere temporal clusters comprising channels 4, 5, 6 and 7 revealed N versus baseline significance from 4.5. –9.25 seconds PST (Figure 4). Similarly patterns were found for HbR, with right hemispheric temporal clusters driving significance (channels 13 – 18 inclusive) from 6.75 – 12.5 seconds PST, while the same temporal cluster as found for HbO (4, 5, 6, 7) drove left hemispheric activation from 6.5 – 11.75 seconds. For V versus baseline, significant activity was lower, with a right hemispheric temporal (channels 15, 16 and 18) driving HbO significance from 4.5 – 9 seconds PST, while temporal channel 9 was the only channel driving significance in the left hemisphere (4.5 – 10 seconds PST). For HbR, a posterior temporal right hemispheric cluster (channels 15 and 18) drove significance from 7 – 13.75 seconds PST. These responses were broadly similar to the GM cohort, though evident in a narrowed regional area in each hemisphere

When contrasting N and V conditions, significant activation in HbO was driven by right hemispheric temporal clusters (channels 13 - 16) from stimulus onset to 5.25 seconds after, with only one channel (4) active for HbO in the left (6 – 6.5 seconds PST). For HbR, activity was more widespread across temporal regions (channels 13-18) in the right hemisphere and located to more medial temporal channels (4, 5, 7 and 9) in the left hemisphere (right time window: 3 – 12.5 seconds PST; left median time window: 6-14.5 seconds PST). For all this activation, N was greater than V, with V<N not significant for any contrasts.

Subsequently, for cross-paradigm analyses the ROI identified in N>V was utilized, with the time window of 5-9 seconds post-stimulus onset used to extract HbO while 7-11 seconds post-stimulus onset was used for HbR for the UK cohort.

#### 3.1.2 HaND

##### GM Cohort

*Habituation*: when contrasting Fam1 versus baseline, TFCE revealed significant activation in HbO driven by a left hemispheric frontal-temporal cluster (channels 1, 4 and 7) and a singular right hemispheric channel (13) starting 5.75 seconds PST and ending 10.5 seconds PST (Figure 6 below). For HbR, a frontal-temporal channel pair (4 and 7) in the left hemisphere drove activation, while the right medial channel 13 drove significance from 5.5 – 12.5 seconds PST. When contrasting Fam3 with baseline, no channels drove HbO significance, while for HbR, left medial-temporal clusters (channels 4, 6 and 9), and right medial-temporal channels (13, 16 and 18) drove significance from 8.75 – 14.5 seconds PST.

**Figure 6.**
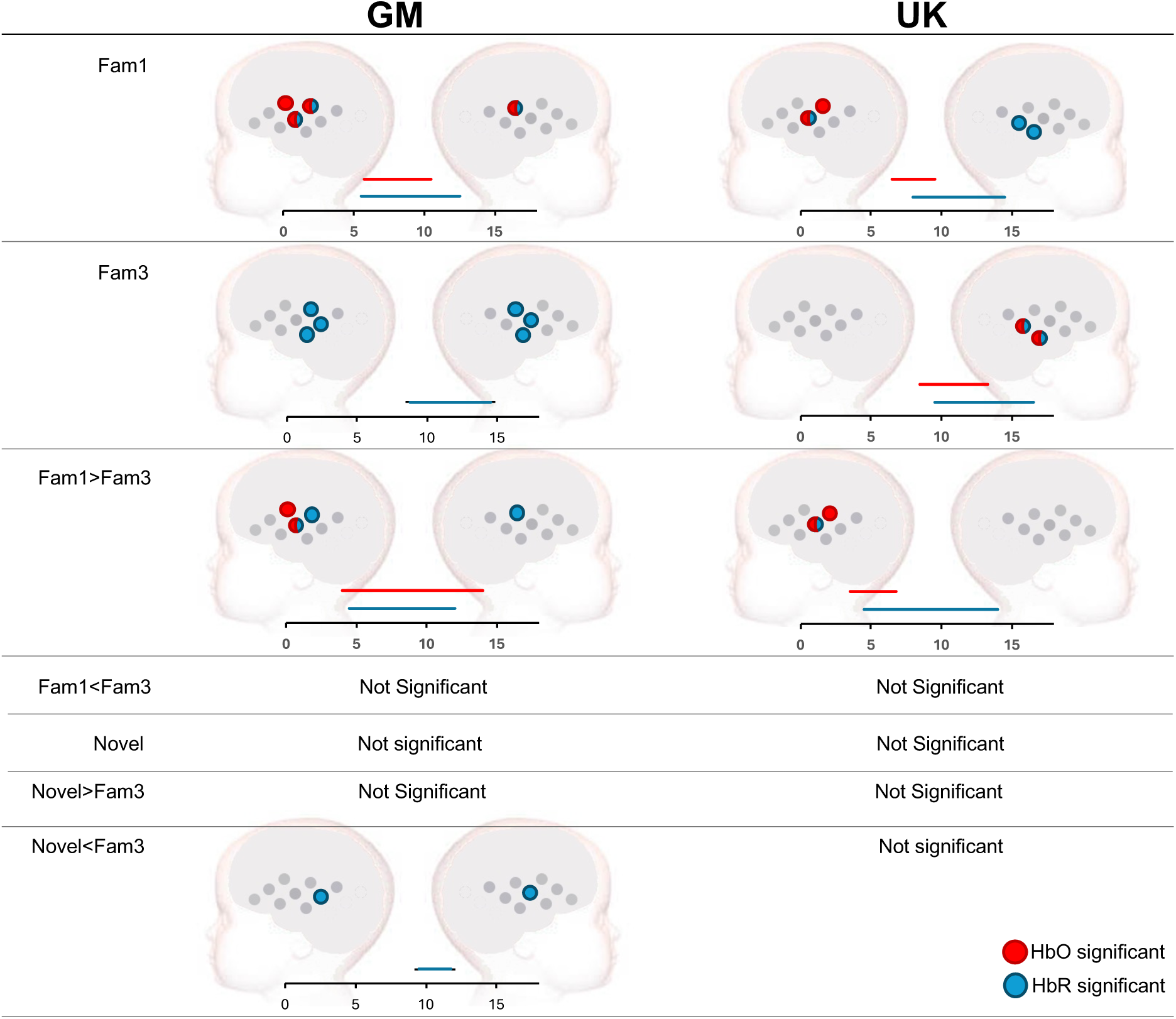
Significant channel by channel haemodynamic activation in the HaND paradigm across GM (left) and UK (right) cohorts. Red indicates channels with significant HbO, blue those with significant HbR. A time-bar has also been included, which indicates when significant activity for HbO and HbR was found for each condition contrast (in sec from stimulus onset).

When contrasting Fam1 versus Fam3, a frontal-temporal cluster on the left hemisphere (channels 1 and 7) drove significant habituation in HbO from 4 – 14 seconds PST. For HbR, significance was driven by a left temporal cluster (channels 4 and 7), and by a singular temporal channel (channel 13), from 4.5 – 12 seconds PST.

*Novelty*: when contrasting Novel versus baseline, no significant contrasts were found for HbO or HbR (Figure 6). For this cohort, two channels (6 and 16) displayed HbR significance in the opposite direction to Novelty (Fam3 > Nov), from 9.5 to 11.5 seconds PST, showing continued habituation to the auditory stimuli regardless of the change in stimuli.

##### UK Cohort

*Habituation*: when contrasting Fam1 versus baseline, TFCE revealed significant activation in HbO across a left hemispheric temporal cluster (channels 4 and 7), driven by activation from 6.5 - 9.5 seconds PST (Figure 6 below). For HbR activity was driven by channel 7 in the left hemisphere, and temporal clustered channels (15 and 18) in the right hemisphere, from 8 – 14.5 seconds PST. When contrasting Fam3 with baseline, clustered right temporal channels 15 and 18 drove HbO significance from 8.5 – 13.25 seconds PST, while the same channels drove HbR significance from 9.5 – 16.5 seconds PST. No channels were identified that continued to show significant changes during Fam3 in the left hemisphere.

When subsequently contrasting Fam1 versus Fam3, left temporal channels (4 and 7) drove significant habituation in HbO from 3.5 – 6.75 seconds PST. For HbR, significance was driven by a singular left temporal cluster (channel 7) from 4.5 – 14 seconds PST. This habituation was in a similar location to the GM cohort in the left hemisphere, however in contrast to the GM cohort significant habituation was not found in the right hemisphere.

*Novelty*: similarly to the GM cohort, no significant Novel versus baseline was found for HbO or and no significance was found when contrasting Novel with Fam3 for this cohort.

Given these results, the identified ROI’s with channels showing significant activation to Fam1> baseline in both the UK and the GM cohorts, were then extracted for HbO (channels 4 and 7) and HbR (channel 7), using the following time windows: 6-10 seconds post-stimulus onset (HbO), and 8-12 seconds post-stimulus onset (HbR), in order to best capture this data across cohorts. Once again, only Habituation data was extracted, given the lack of a Novelty response in either cohort.

When data was averaged across all channels with significant responses within epoch(s) and participants, Habituation can be seen due to the larger HRF during Fam1 compared with Fam3 (Figure 5).

#### 3.1.3 Functional Connectivity GM Cohort

Infants demonstrated stronger inter-hemispheric connectivity compared to intra-hemispheric connectivity (for HbO: average inter-hemispheric connectivity R value across the different inter-hemispheric ROI pairs averaged across all infants = 0.613, average intra-hemispheric R = 0.331; for HbR: average inter-hemispheric R = 0.540, average intra-hemispheric R = 0.376). A paired-samples t-test revealed a significant difference between inter-and intra-hemispheric connectivity (HbO: t_(120)_=17.748, p<0.001; HbR: t_(120)_=12.357, p<0.001). As shown in Figure 7, the majority of individuals in the GM cohort show inter-connectivity to be higher than intra-connectivity.

**Figure 7.**
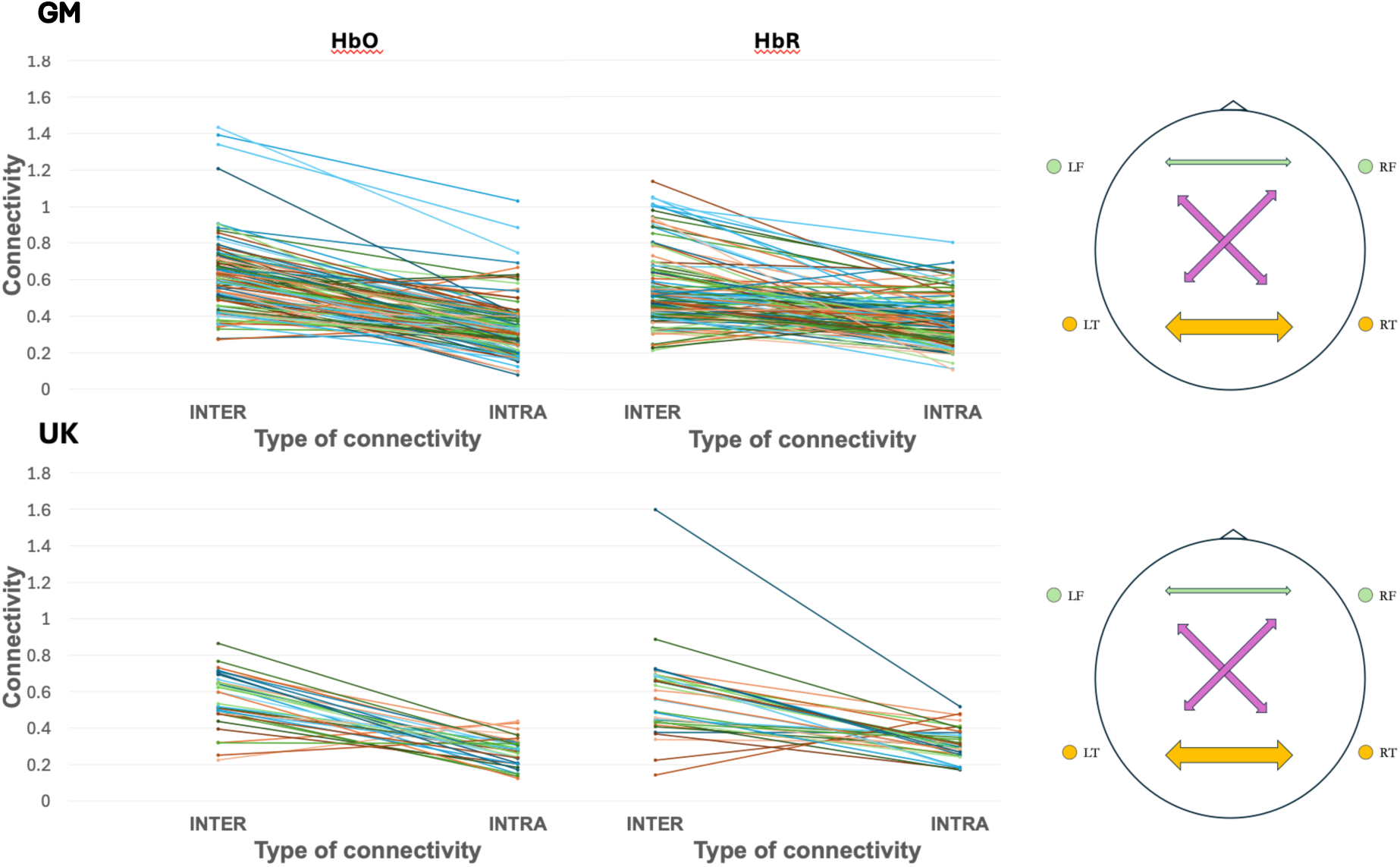
Inter-and intra-hemispheric connectivity for Gambian (top) and UK (bottom) infants. In the left (HbO) and middle (HbR) panels the overall grand averaged inter-and intra-hemispheric connectivity values are displayed for each individual infant (one line per infant). In the right panel, a visual representation of the approximate strength of regionally specific inter-hemispheric connections are depicted (with the largest weighted arrows indicating large R values (> 0.62), the medium weighted arrows indicating medium R values (0.5 – 0.62) and the smallest weighted arrows indicating low R values (<0.1); green – frontal-frontal; pink – frontal-temporal; orange – temporal-temporal).

When examining which connections drove this stronger inter-hemispheric connectivity, temporal inter-hemispheric connectivity had the strongest correlation (HbO: R=0.762, HbR: 0.699), while between region inter-hemispheric connectivity moderately correlated (left frontal, right temporal: HbO: R = 0.528, HbR: 0.521, right frontal, left temporal: HbO: R = 0.506, HbR: 0.532). In contrast, frontal inter-hemispheric connectivity had a weak correlation (HbO: R=0.012, HbR: 0.025) (Figure 7).

Repeated measures ANOVA revealed a significant effect of region on inter-hemispheric connectivity, such that temporal inter-hemispheric connectivity was significantly stronger than frontal and between-region inter-hemispheric connectivity. Significance was maintained upon corrected degrees of freedom (using Greenhouse-Geisser, given assumptions of sphericity were violated (F_(120, 3)_=23.63, p<0.0001)), with region accounting for a moderate proportion of the variance in within-subject inter-hemispheric connectivity (η²_G = 0.12). Post-hoc tests (FDR corrected for multiple-comparisons) revealed significant differences between temporal inter-hemispheric connectivity (IHC) and: (i) frontal IHC (t_(120, 3)_=8.286, p<0.0001), (ii) right frontal-left temporal IHC (t_(120, 3)_=3.586, p<0.001), and (iii) left frontal - right temporal IHC (t_(120, 3)_=2.854, p<0.01). HbR results confirmed the same patters on the repeated measures ANOVA (F_(120, 3)_=4.70, p<0.0001, η²_G = 0.10) and FDR corrected post-hoc tests contrasting inter-temporal connectivity with frontal IHC (t_(120, 3)_=7.218, p<0.0001), right frontal-left temporal IHC (t_(120, 3)_=2.812, p<0.01), and left frontal - right temporal IHC (t_(120, 3)_=2.232, p<0.05).

Given the significance of inter-hemispheric connections at both the total and regional level, these correlations were taken forward for cross-paradigm analyses.

### UK Cohort

The same overall patterns observed in Gambian infants, were found in the UK, with stronger inter-hemispheric connectivity than intra-hemispheric connectivity reported on average (for HbO: inter-hemispheric R = 0.562, average intra-hemispheric R = 0.272; for HbR: inter-hemispheric R = 0.577, intra-hemispheric R = 0.314). A paired-samples t-test revealed a significant difference between inter-and intra-hemispheric connectivity (for HbO: t_(40)_=9.271, p<0.001, HbR: t_(40)_=6.563, p<0.001). As shown in Figure 7, the majority of individuals in the UK show inter-connectivity to be higher than intra-connectivity.

When examining which connections drove stronger inter-hemispheric connectivity, temporal IHC had the strongest correlation (HbO: R=0.748, HbR: 0.636), while between regions IHC moderately correlated (left frontal, right temporal: HbO: R = 0.550, HbR: 0.606, right frontal, left temporal HbO: R = 0.590, HbR: 0.618), and frontal IHC was weak (HbO: R = 0.007, HbR: 0.086) (Figure 7).

A repeated measures ANOVA using corrected degrees of freedom (Greenhouse-Geisser) found a significant difference across inter-hemispheric connections for HbO (*F*(31,3) = 10.321, *p* <.0001), with between-region differences accounting for a moderate amount of the within-participant variance (η²_G = 0.20). Post-hoc pairwise comparisons, FDR-corrected for multiple testing, indicated that this effect was primarily driven by significantly greater inter-temporal connectivity compared to inter-frontal connectivity (*t*(31,3) = 5.94, *p* <.0001) and right frontal - left temporal ( *t*(31,3) = 2.359, *p* <.05). Differences between inter-temporal and left frontal–right temporal connections did not reach significance.

HbR results confirmed the same patterns on the repeated measures ANOVA (F_(31, 3)_=1.813 p<0.0001, η²_G = 0.031) although FDR corrected post-hoc tests, found no significant contrasts between inter-temporal connectivity and IHC between other regions.

As in the GM cohort, the significance of inter-hemispheric connections at both the total and regional levels meant these correlations were taken forward for cross-paradigm analyses.

### 3.2 Cross-Paradigm Analyses

#### 3.2.1 Functional Activation Paradigms

##### Is condition selectivity in the *Social* paradigm correlated with habituation to repeating condition trials in the *HaND* paradigm?

To examine for condition selectivity, contrasts were extracted for each paradigm examining: associations between non-vocal versus vocal responses (Social paradigm) and Fam1 versus Fam3 (HaND paradigm). No significant correlations were found between Habituation and non-vocal / vocal selectivity, for either cohort.

#### 3.2.2 Functional Activation and Functional Connectivity

##### Do HaND or Social paradigm responses relate to functional connectivity?

In the HaND paradigm, no associations were found between total interhemispheric connectivity and habituation for either cohort. However, given the significant difference between temporal and frontal interhemispheric connectivity within ROIs this was further explored at the region level. Temporal IHC negatively correlated with habituation on the HaND paradigm, post Bonferroni correction (R =-0.197,_df=108_, p=0.0415) for the GM, but not UK (r = 0.017, p = 0.9271, n = 31), cohort (Figure 8). While this association suggests the strength of temporal interhemispheric FC may relate to whether infants dishabituate during sleep (39.8%) or habituate (60.2%) across repeating trials in the HaND paradigm, these results are borderline significant with a low effect size and should therefore be taken with caution. No other associations were found to be significant.

**Figure 8.**
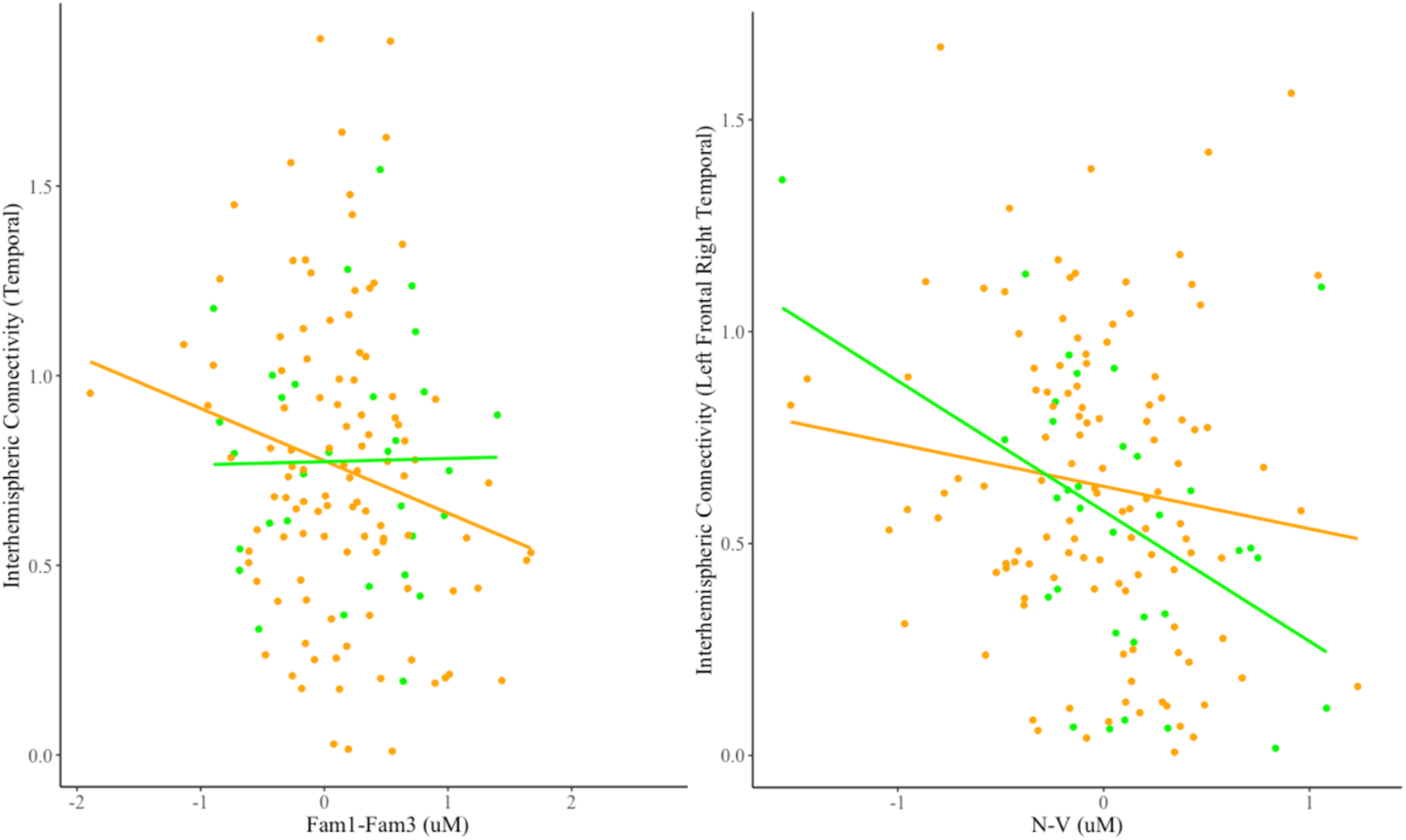
Scatterplots showing the correlation between the functional paradigm HbO responses and HbO interhemispheric connectivity (y-axis), with lines of best fit for the GM (orange) and UK (green) cohorts. The left panel shows the total inter-hemispheric connectivity and the averaged habituation response (Fam1 > Fam3), with a positive value on the x-axis indicating higher habituation across trials. The middle panel shows the total inter-hemispheric connectivity and strength of non-vocal selectivity (non-vocal > vocal contrast) with a negative value on the x-axis indicating higher social selectivity and a positive indicating higher non-social selectivity. The right panel shows the regionally specific inter-hemispheric connectivity between the left frontal and right temporal regions and strength of non-vocal selectivity (non-vocal > vocal contrast).

In the *Social Selectivity* paradigm, no associations were found between total interhemispheric connectivity and selectivity for either cohort. However, as with the *HaND* paradigm, given the association between temporal and frontal interhemispheric connectivity this was further explored at the region level. When examining the *Social Selectivity* paradigm, significant associations were found in the UK cohort, and trending towards significance in the GM cohort. Contrasting left frontal - right temporal inter-hemispheric connectivity with non-vocal selectivity (N-V) revealed a significant negative correlation (R=-0.453, _df=33_, p<0.008), of medium effect size, in the UK cohort (Figure 8), showing that those infants with stronger vocal selectivity had stronger overall inter-hemispheric connectivity. In the GM the same negative association could be seen (Figure 8), however this did not reach significance (r =-0.132, p = 0.156, n = 117). No other associations were found to be significant.

## 4.0 Discussion

The present study examined early infant brain imaging data from two contrasting populations (Gambia and UK) across three different neuroimaging paradigms to better understand early markers of neonatal neurodevelopment. This work additionally aimed to ascertain whether cross-paradigm correlates can provide more insights into infant brain maturation, and to aid understanding of whether differences in such developmental markers vary across settings.

Within individual paradigms, infants across The Gambia and UK had some overlapping responses: (i) both cohorts demonstrated significant habituation to an auditory *HaND paradigm*, with responses strongest in the left hemispheric temporal regions, (ii) neither cohort were found to have significant novelty detection, (iii) for the *Social selectivity paradigm* infants demonstrated relatively widespread, bilateral activation in response to non-vocal stimuli, while vocal reponses were more robustly lateralized to the right hemisphere (a greater number of channels activated in the right compared with the left hemisphere for vocal stimuli with both HbO and HbR significant), and (iv) for the *Functional Connectivity paradigm* inter-hemispheric connectivity was stronger than intra-hemispheric connectivity at the group level, with this particular result driven by temporal compared to frontal, inter-hemispheric connections. While a number of between-cohort similarities were evident at one-month of age, two main differences were noted: (1) Gambian infants did not demonstrate any significant contrast to non-vocal versus vocal stimuli in the Social paradigm at group level, while UK infants reported non-vocal selectivity; and (2) Gambian infants continued to habituate across novelty trials at the group level suggesting a change in stimulus was not detected.

For the cross paradigm analyses there were two main findings. Firstly, no associations were found between the functional paradigms, revealing that social auditory selectivity to individual sounds did not associate with patterns of habituation to repeating speech in longer form sentences at one month of age. This supports the hypothesis that individual responses across these paradigms are distinct, are not interrogating similar cognitive processes and are therefore each able to identify a unique biomarker of early development. Secondly, left frontal - right temporal inter-hemispheric connectivity associated with social selectivity (in the UK, and trending in the Gambia cohort), showing that those infants with stronger vocal selectivity had stronger overall inter-hemispheric connectivity. This supports the hypothesis that those infants with more mature connectivity profiles show more rapid specialisation of social discrimination responses.

One-to-five-day old infants have been found to display widespread neural habituation to repeated auditory stimuli, while by 5-months of age this effect has been localised to left hemispheric temporal regions^19,65^. Results from the present study indicate that such left hemispheric localisation could emerge within the first month of life, and that this may be consistent across infants from two very diverse global populations. In contrast to habituation, significant novelty responses were not found in either cohort in the *HaND* paradigm at this age point. In analyses of longitudinal profiles of the HaND responses of this same BRIGHT cohort at later age points (5, 8, 12, 18 months, 2 and 3-5 years), novelty was reported to begin to emerge by 5-months of age in UK infants, although across the first years of life this development was nonlinear^19,22^. In contrast, in Gambian infants novelty detection emerged by 18-24 months of age^22^, and in line with the current findings, infants continued to habituate across novelty trials up to 12 months of age. This could indicate that, while similar developmental markers in HaND are apparent at one-month of age, and demonstrate early localisation of habituation, discrepancies may begin to emerge and strengthen across the first year of life.

While patterns of habituation and novelty detection were broadly similar across cohorts at one-month of age, group responses to the *Social Selectivity* paradigm were already notably different. In line with previous findings that non-vocal selectivity precedes vocal^54,66,67^, infants in the UK demonstrated non-vocal selectivity. Gambian infants did not show this response at group level, which is surprising given that in a previous study in a different cohort of one month old Gambian infants non-social selectivity was evident^10,54^. During pregnancy, procesing of prosodic sounds is supported by a functionally active but relatively immature, posterior auditory network that operates via bottom-up processing^68^. Postnatally, auditory stimuli take on a new intensity and complexity, and the auditory network transitions to a frontal-temporal, top-down network which facilitates enhanced encoding and decoding of auditory stimuli^68^. Interestingly, the cross-paradigm analyses revealed a high degree of individual variability in the social-selective response, with some infants evidencing vocal>non-vocal selectivity and others showing non-vocal>vocal selectivity across both cohorts. Therefore, the different group level findings seen across different studies could reflect the high degree of individual variance evident in the first months of life, and by extension, that vocal selectivity is an early marker of brain maturity. In a later section we discuss how this is further supported by the cross-paradigm analyses.

Early life brain responses to auditory stimuli and language, have been shown to be heavily dependent on environmental exposures to auditory stimuli such as conversational turns as well as to nonvocal and nonsocial noise, with subsequent impacts upon the development of stimuli-related responses as well as underlying interhemispheric connectivity^42,43,68^. When considering such cross-paradigm findings, these results could reflect the different home exposures more likely experienced across these two cultures. In the Gambian West Kiang region, the average family unit is comprised of 16 members, with exposure to a higher density and diversity of auditory stimuli more probable given the proximity of many people living together^49^. In contrast, most UK participants were from middle-class families that were typically comprised of two adults, which may limit social and auditory exposures more heavily towards dense interactions with a lower number of social partners in the first month of life. While cohort differences were not statistically examined in the present study, disparities in these home environments could, understandably, yield different brain responses across paradigms. However, further research that statistically examines cohort differences and incorporates wider environmental variables would be needed to better untangle this.

Previous research into functional connectivity has indicated that intra-hemispheric connections precede the growth of inter-hemispheric connections in early development, with short-range, intra-hemispheric connectivity more prevalent prenatally and thought to support local specialisation of brain regions. Postnatally, intra-hemispheric connections decrease via synaptic pruning, while longer range inter-hemispheric connections develop, helping to integrate more global networks over the first years of postnatal life^20,42,43^. Given that inter-hemispheric connectivity is posited to be a marker of infant age or brain maturity in early development, the present study’s findings across both Gambian and UK cohorts, align with this early postnatal transition. Such connectivity appeared to be primarily driven by temporal connectivity when compared with frontal and inter-regional connections, with this holding true across cohorts.

Furthermore, the strongest inter-regional connectivity was found to be between right temporal and left frontal regions. In addition, social selectivity was found to associate with the individual variance in inter-hemispheric right temporal to left frontal connectivity patterns. Compellingly, rapid maturation in left frontal to right hemisphere connectivity has been previously reported as being associated with auditory network transitions to a frontal-temporal, top-down network^68^.

Some limitations need to be acknowledged. While we explored similarities and differences between responses across infant cohorts as potentially being driven by individual differences in developmental specialisation and maturity of brain connectivity, as well as the result of environmental influence, several methodological factors could have underlied these findings as well. Discrepancies between study results across our cohorts, and relative to previous findings, could be due to changes in fNIRS preprocessing and analysis pipelines, or differences in sample size. Furthermore, given all paradigms were undertaken during infant sleep, the associations (or lack of) found could be influenced by infant sleep stages. This is especially the case given that paradigms were conducted in the same order at each presentation, which may alter the likelihood of certain sleep stages being linked to particular paradigms and which, in turn, can impact the functional brain networks activated^43,69,70^. While examination of sleep stages went beyond the scope of the present study, a separate paper analysing the impact of sleep stages on paradigm responses explores this factor^71^. A second limitation is that the UK sample was much smaller than the Gambian, resulting in lower power of statistical analyses in this cohort. Thirdly, functional connectivity data was analysed using a slightly different pre-processing pipeline, due to the nature of FC data, which could have impacted the comparability results. Finally, much of this research was exploratory, given its novelty.

The present study had a number of strengths including the inclusion of two cohorts of infants, from contrasting global populations and settings, but using identical testing protocols, paradigms and analyses; its use of threshold-free cluster enhancement to identify regions of interest and time-windows; its use of cross-paradigm analyses that utilized comparable pre-processing pipelines, and its examination of infant outcomes at the very early time-point of one-month of age. The present study also highlighted measures upon which infant outcomes may be comparable across settings, as well as those areas where early developmental trajectories may already diverge. Important takeaways about early infant developmental trajectories and markers of maturation across contexts were identified.

## 5.0 Conclusions

Overall the current study determined that: (i) more robust habituation and greater vocal selectivity (UK) as well as higher rates of inter-hemispheric and integrated regional connectivity (UK and GM) may represent more mature brain development in the first months of life, (ii) use of these markers together may give deeper and more robust insights into neurodevelopmental trajectories compared to examining each marker alone, (iii) Gambian and UK infants already demonstrate some divergence in their neurodevelopmental trajectories at one-month of age, and (iv) these differences may highlight more optimal or pertinent markers of neurodevelopment in infants from different global settings. Future research could benefit from repeating the same analyses at later time points in the BRIGHT Project to understand whether the identified markers across cohorts develop longitudinally both within-and across-paradigms. Moreover, the inclusion of contextual variables to identify those most pertinent environmental exposures across, or between, settings would help improve understanding of whether differences across cohorts are optimally adaptive or, in some instances, may reflect the early emergence of developmental delay or diversity. Nonetheless this study is one of the first to demonstrate task-specific response patterns and group level differences in an extensive neuroimaging battery during the neonatal period, holding the potential to inform the future use of these markers in early identification and intervention.

## Notes

**Acknowledgements:** We are indebted to the parents and infants who took part in this project without whom this work would not have been possible. We also thank the wider team of staff at the MRC Keneba Field Station for supporting us in the collection of this data. The BRIGHT Project was funded by the Gates Foundation (OPP1127625) and core funding (MC-A760-5QX00) to the International Nutrition Group by the Medical Research Council UK and the UK Department for International Development (DfID) under the MRC/DfID Concordat agreement. This work was further supported by the Economic and Social Research Council, Doctoral Training Pathway Studentship to IG; the Medical Research Council Programme Grant (MR/T003057/1) to BB and the UKRI Future Leaders fellowship (MRC grant MR/S018425/1) to SLF. The views expressed are those of the authors and not necessarily those of the GF, ESRC, MRC or the UKRI.

### Competing Interest Statement

The authors have declared no competing interest.

