## Supplementary information for "Cross-Paradigm fNIRS Brain Activity in Neonates across The Gambia and UK"

Supplemental Materials: ***S1. Cross-Paradigm Results with HbR***

At one-month of age HbO and HbR responses are attenuated compared to later childhood and adulthood. This can make deciphering responses more challenging, with HbO providing a more robust paradigm-related response (amplitude) in comparison to HbR given it's more favourable signal-to-noise ratio (Lloyd-Fox et al., 2010; Gervain et al., 2011). For cross-paradigm analyses within the main paper, HbO was therefore extracted and taken forward. However, given that HbR has been posited to have heightened accuracy compared to HbO, both chromophores were examined, with HbR reported here in supplementary analyses using the same methods as employed for HbO.

| | **TOTAL Inter** | **Frontal** | **Temporal** | **L-F to R-T** | **R-F to L-T** |
|---|---|---|---|---|---|
| | | | **UK** | | |
| **Habituation (HaND)** | -0.161 | -0.075 | -0.095 | 0.036 | -0.14 |
| **Nonvocal-vocal** | 0.122 | 0.204 | 0.098 | 0.195 | 0.209 |
| | | | **GM** | | |
| **Habituation (HaND)** | -0.084 | 0.017 | -0.016 | 0.058 | 0.087 |
| **Nonvocal-vocal** | -0.049 | 0.013 | 0.04 | -0.083 | -0.017 |

**Table 1.** Cells represent correlation coefficients (R values) for defined variables in the x and y columns. Mean Inter = mean total interhemispheric connectivity; Frontal = mean frontal interhemispheric connectivity; Temporal = mean temporal interhemispheric connectivity; L-Frontal R-Temporal = Left Frontal – Right Temporal interhemispheric connectivity; R-Frontal L-Temporal = Right Frontal – Left Temporal interhemispheric connectivity. No correlations reached significance at the $p<0.05$ level. Moreover, for GM Mean Inter, Inter Frontal and Left Frontal - Right Temporal values were log transformed, given non-normality, while for UK Mean Inter and Right Frontal – Left Temporal values were log transformed.

No significant correlations were found between HaND or Social data and IHC for HbR. Differences in response amplitude; ROI, or time windows used for HbO and HbR could underlie such discrepancies between HbO and HbR findings. However, further research to better untangle the associations of functional brain activation responses across paradigms and with functional connectivity, would help better understand discrepancies seen at the one-month age point.
